# Edible Plant-Derived Extracellular Vesicles Preserve Secondary Metabolites and Exhibit Source-Dependent Biological Activities

**DOI:** 10.64898/2026.08.17.742983

**Authors:** P. Debishree Subudhi, Vibhuti Jakhmola, Shruti Chalil Sureshan, VengalaRao Yenuganti, Neha Saroj, Shivani Gautam, Prachi Sinha, Chhagan Bihari, Shiv Kumar Sarin, Sukriti Baweja

## Abstract

Edible plant foods are rich sources of bioactive secondary metabolites that contribute to various health beneficial properties. Naturally occurring plant-derived extracellular vesicles (PDEVs) may serve as food nanocarriers that protect and facilitate the oral delivery of these metabolites; however, their source-specific composition, gastrointestinal stability, and functional properties remain completely unknown. In this study, PDEVs were isolated and comprehensively characterized from four phytochemically distinct edible plant foods: black carrot, ginger, garlic, and turmeric. Gastrointestinal stability was evaluated in a simulated digestion model. Untargeted LC–MS/MS metabolomics identified 572 metabolites with distinct source-specific signatures, including lignans and quercetin derivatives in black carrot EVs, [6]-gingerol and silymarin in ginger EVs, diosgenin in garlic EVs, and curcumin in turmeric EVs. Pathway analysis associated these metabolites with antioxidant, anti-inflammatory, epithelial barrier, lipid metabolic, and apoptosis-related functions. Functionally, carrot EVs enhanced claudin and occludin expression, while ginger EVs restored ZO-1 and reduced cyclin D1 and MMP9 in ammonia-stressed intestinal epithelial cells. Garlic and turmeric EVs suppressed STAT3, AKT1, and TNF-α, whereas garlic EVs further attenuated steatosis by decreasing PNPLA3 and SREBP-1c in steatotic hepatocytes. These findings demonstrate that edible PDEVs are gastro-intestinally stable nanocarriers of bioactive secondary metabolites with source-specific functional properties, supporting their potential as functional food ingredients and nutraceuticals.

**Graphical abstract:** 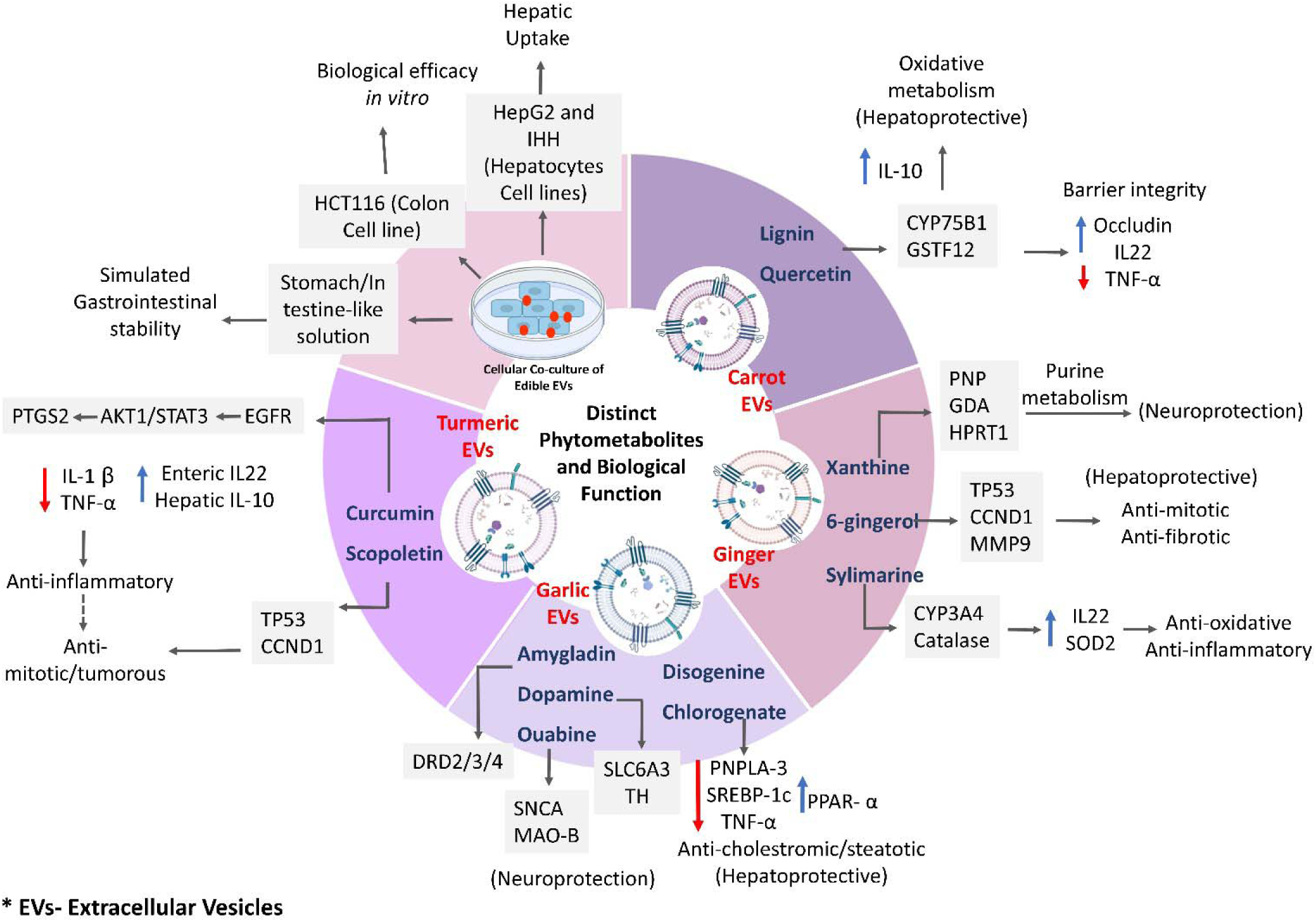

## Introduction

Edible plant foods are rich reservoirs of structurally diverse secondary metabolites, including polyphenols, flavonoids, anthocyanins, terpenoids, alkaloids, curcuminoids, and organosulfur compounds, that contribute to their antioxidant, anti-inflammatory and antimicrobial properties, which has various health benefits [1]. These phytochemicals have attracted considerable attention as functional food ingredients and nutraceuticals owing to their ability to modulate multiple biological pathways associated with chronic diseases [2]. However, many dietary phytometabolites exhibit poor gastrointestinal stability, limited bioavailability, rapid metabolism, and inefficient cellular uptake, thereby restricting their biological efficacy following oral consumption [3]. Consequently, identifying naturally occurring food matrices capable of protecting and facilitating the delivery of these bioactive compounds has become an important focus in food science and nutrition research [3, 4].

Edible plant-derived extracellular vesicles (PDEVs) are nanosized lipid bilayer vesicles naturally released by fruits, vegetables, and medicinal plants [5]. Unlike conventional plant extracts, PDEVs encapsulate diverse biomolecules, including proteins, lipids, nucleic acids, and metabolites, within a protective membrane that enhances their stability and facilitates cellular uptake [6, 7]. Because they originate from commonly consumed edible plants, PDEVs are considered inherently biocompatible, exhibit excellent gastrointestinal stability, and have emerged as naturally occurring food nanostructures capable of preserving and transporting dietary phytochemicals. These unique characteristics have generated growing interest in exploiting PDEVs as functional food ingredients and nutraceutical delivery systems [8].

Increasing evidence indicates that edible PDEVs possess intrinsic biological activities in addition to serving as carriers of bioactive molecules. PDEVs isolated from ginger, grapefruit, lemon, grape, carrot, and aloe vera have demonstrated antioxidant, anti-inflammatory, intestinal barrier-protective, hepatoprotective, antimicrobial, and immunomodulatory properties in various experimental models[9–13]. Nevertheless, these studies have largely investigated individual plant species using different isolation procedures, characterization strategies, and experimental models, making direct comparisons difficult and limiting our understanding of how differences in phytochemical composition influence biological function [5, 8].

Although proteins, lipids, and small RNAs have been identified in edible PDEVs, whereas, there is no information available on their associated metabolites. Since secondary metabolites are the principal bioactive constituents responsible for the health-promoting properties of edible plants, comprehensive comparative characterization of PDEV-associated metabolites is essential for understanding their nutritional significance. Moreover, little is known about how phytometabolite composition varies among edible plant sources or how these differences influence gastrointestinal stability and source-specific biological activities. In addition, inconsistent physicochemical characterization and limited adoption of the Minimal Information for Studies of Extracellular Vesicles (MISEV) guidelines continue to hinder reproducibility and cross-study comparisons in this rapidly expanding field [14].

To address these knowledge gaps, we selected four phytochemically distinct edible plant foods black carrot (*Daucus carota* L.), ginger (*Zingiber officinale*), garlic (*Allium sativum*), and turmeric (*Curcuma longa*) representing diverse classes of health promoting secondary metabolites, including anthocyanins, gingerols, organosulfur compounds, and curcuminoids, respectively. We hypothesized that the phytometabolite composition of edible PDEV is source-dependent and contributes to their gastrointestinal stability and functional properties. Therefore, we performed a comprehensive comparative characterization of PDEVs using transmission electron microscopy, nanoparticle tracking analysis, zeta potential measurements, simulated gastrointestinal digestion, and high-resolution untargeted LC–MS/MS metabolomics, followed by pathway enrichment and functional validation in intestinal epithelial and steatotic hepatocyte models. By linking source-specific metabolite signatures with biological activities, this study provides new insights into edible plant-derived nanovesicles as naturally occurring carriers of food bioactive compounds and supports their potential application as functional food ingredients and nutraceuticals for promoting various health benefits.

## Methods

### Isolation of Edible Extracellular vesicles

Black carrot (*Daucus Carota*; Pusa Asita, IPC-126; notified vide S.O. 3666(A), 6 Dec 2016), ginger (*Zingiber officinale*; KGWF-01/23), garlic (*Allium sativum*; IC No. 375117) and turmeric (*Curcuma longa*; KTFF-01/23) were obtained from accredited horticultural institutions. Edible parts of the plants from various sources were peeled, thoroughly washed, and processed into extract under aseptic conditions. The initial procedures with separation of the debris and cellular remnants through sequential centrifugation performed at 1,000 × g for 10 min and 3,000 × g for 20 min, followed by isolation of edible EVs from the clarified supernatant using sucrose density gradient ultracentrifugation at 100,000 × g for 120 min (Schematic representation in Suppl. Fig. S1), as described previously [8]. Of the density gradient ultracentrifugation fractions corresponding to the 30%/45% sucrose interface were collected and resuspended in PBS for further analysis. The final EV pellet was resuspended in sterile, double-filtered 1× PBS and stored at 4 °C for immediate use or −80 °C for long-term storage.

### Nanoparticle Tracking Assay and Zeta Potential Measurement

Purity and vesicular integrity were assessed through nanoparticle tracking analysis (NTA) and zeta potential measurement. The hydrodynamic size and particle concentration of edible EVs were determined by NTA using a NanoSight LM20 system (Malvern Panalytical, UK) equipped with a 405 nm laser and a scientific CMOS camera. Data were acquired and analyzed using NTA software version 3.1. For each sample, three 60 s videos were recorded at room temperature (Suppl. Video), with the camera level and detection threshold kept constant across all measurements to ensure comparability. Particle size distribution profiles were calculated, and the mean particle diameter (nm) and concentration (particles/mL) were reported. All samples were diluted in 0.22 μm-filtered PBS to achieve a particle count within the optimal detection range (30–100 particles/frame).

The surface charge (ζ-potential) of edible EVs was measured using Zetasizer Ultra (Malvern Panalytical Ltd., Malvern, UK) based on electrophoretic light scattering (ELS). Samples were analyzed in disposable folded capillary cells (DTS1070) at 25 °C, and the electrophoretic mobility was converted to zeta potential using the Smoluchowski approximation. Instrument calibration and quality control were performed according to the manufacturer’s guidelines prior to each session. These distinct and complementary characterization approaches provide strong evidence that the isolated particles represent bona fide edible EVs rather than non-vesicular contaminants. The reproducibility of these features across independent biological replicates and plant batches further supports the reliability of the isolation procedure.

### Transmission Electron Microscopy

The morphology of edible EVs was analyzed using a Talos F200C transmission electron microscope (Thermo Fisher Scientific, 200 kV). Freshly isolated EVs were loaded on formvar/carbon-coated copper grids for 5 minutes, rinsed gently with decarbonated distilled water to remove unbound particles, and air-dried. Imaging was performed at low dose to preserve vesicle structure. The obtained micrographs showed round, bilayer vesicles with typical EV morphology, consistent with the MISEV 2024 guidelines [14]. The combined characterization data from TEM, NTA, and zeta potential analyses confirmed vesicle integrity and purity, supporting compliance with MISEV recommendations.

### Stability and Viability of edible Extracellular Vesicles

The stability of edible EVs under gastrointestinal conditions was evaluated using simulated stomach-like and intestine-like solutions, following a protocol similar to INFOGEST [15], with modifications to suit EDIBLE EVS administration. The oral phase was omitted, as edible EVs were not subjected to mastication. For the stomach simulation, edible EVs were incubated in simulated gastric fluid (SLS) containing pepsin (Sigma-Aldrich, 1 mg/mL) and 0.1 M HCl, adjusted to pH 2.0, for 2 hours at 37 °C with gentle agitation using a magnetic stirrer to mimic gastric peristalsis. Following this, the edible EVs were transferred to simulated intestinal fluid (ILS) composed of pancreatin (Sigma-Aldrich, 1 mg/mL) and 70 µl exosome-free bile that was prepared by ultracentrifugation at 100,000 × g for 2 hours to remove endogenous vesicles, adjusted to pH 6.5, and incubated for 2 hours at 37 °C under gentle agitation. After the sequential incubation, edible EVs were collected, and their structural stability and integrity were assessed using NTA for particle size and a Zeta analyzer.

### Untargeted Metabolomics of Edible Extracellular Vesicles

Edible EVs (10^11^–10^12^ particles/mL) were reconstituted in water and subjected to LC–MS/MS analysis following a modified protocol of Altadill *et al.* [16]. Metabolomic profiling was performed using three independent biological replicates, each derived from separate plant batches, to ensure reproducibility across biological and extraction variability. Metabolite extraction was performed through three freeze–thaw cycles. Samples were vortexed (2–3 min), centrifuged (20,000 rpm, 4 °C, 10 min), and the resulting supernatants were collected and dried under vacuum. Dried extracts were reconstituted in 100 µL of solvent (5% acetonitrile, 5% internal standard, 90% water), of which 80 µL was used for LC–MS/MS analysis, while 20 µL was pooled for quality control. Analyses were performed on a Thermo Scientific™ UHPLC system coupled to a Q Exactive™ Orbitrap mass spectrometer, operating in both positive and negative electrospray ionization modes, with separation achieved on a Hypersil GOLD™ C18 column. The metabolite-protein interactome analysis was performed using the STITCH 5.0 database. Metabolite identities were putatively annotated on the basis of accurate mass and retention time matching against reference spectral and compound databases; no MS/MS-based spectral confirmation or authentic standard verification was performed. These annotations therefore represent compositional signatures rather than structurally confirmed identities and should be interpreted accordingly.

### *In vitro* Co-Cultures and Uptake of Edible EVs

To evaluate the pharmacological activities of edible EVs, HCT116 enterocytes were treated with 20 mM NH₄Cl to induce oxidative and inflammatory stress, followed by co-culture with edible EVs. Their effects were examined on epithelial barrier integrity (ZO-1, occludin), cell proliferation/cycle (p53, cyclin D1 and MMP9), apoptosis (caspase 3), cell signaling (akt 1, stat 3, and ptgs 2), inflammatory mediators (TNF-α, IL-22), and oxidative stress response (cyp1A1).

Isolated edible EVs were labeled with PKH26 dye (Sigma, Mini26-1KT, SLBW0232) following the manufacturer’s protocol. PKH26-labeled edible EVs were co-cultured with HepG2 hepatic cells and analyzed by flow cytometry [17, 18]. For confocal microscopy, PKH26-labeled carrot EVs were co-cultured with immortalized human hepatocytes (IHH) cells, and live-cell imaging was performed with image acquisition every 5 seconds to monitor vesicle uptake. Fluorescence intensity was quantified using ImageJ software.

Hepatoprotective and lipid-regulatory properties were investigated in IHH rendered steatosis by treatment with 0.5 mM BSA-conjugated palmitic and oleic acids. In this model, edible EVs were analyzed for their ability to modulate lipogenic genes (PNPLA3, SREBP-1c, PPAR-α) and inflammatory cytokines (TNF-α, IL-1β, IL-10) through quantitative mRNA expression profiling. RNA was extracted using TRIzol, precipitated with isopropanol, and resuspended in nuclease-free water. After assessing purity, 1 µg of RNA was reverse-transcribed, and quantitative real-time Polymerase Chain Reaction (qRT-PCR) was performed using SYBR Green, with expression normalized to housekeeping genes via the ΔΔCt method. To ensure reproducibility, all assays were conducted in at least three independent biological replicates, and edible EVs were prepared from multiple plant batches. Comparable outcomes were obtained across replicates and batches, indicating consistency of the observed effects.

### Statistical Analysis

Statistical analyses were performed using the Kruskal–Wallis test to evaluate differences among groups. Given the non-parametric nature of the in vitro RT-PCR data and the limited number of biological replicates, no post hoc correction for multiple comparisons was applied. Data are presented as mean ± SD. All statistical analyses and graphical representations were performed using GraphPad Prism version 9.0. Fluorescence intensity was quantified using ImageJ software.

## Results

### Physiochemical Characterization of PDEVs

TEM imaging demonstrated predominantly cup-shaped vesicles, consistent with the typical morphology of extracellular vesicles (Fig.1A). NTA demonstrated heterogeneous (polydisperse) populations of edible EVs (Fig. 1B). The predominant particle sizes (mode) were 60.0 nm for carrot EVs, 197.5 nm for ginger EVs, 214.1 nm for garlic EVs, and 95.8 nm for turmeric EVs. Carrot EVs exhibited a particle size distribution (D10–D90: 56.7–265.4 nm) with a particle concentration of 5.12 × 10^9^ particles/mL. Similarly, ginger EVs displayed a broad particle size distribution (D10–D90: 117.7–392.6 nm) with a particle concentration of 1.17 × 10^9^ particles/mL. Garlic EVs exhibited a particle size distribution ranging from 110.4 to 374.8 nm (D10–D90) and a concentration of 2.68 × 10^9^ particles/mL, whereas turmeric EVs showed a D10–D90 range of 155.7–393.7 nm with a particle concentration of 9.31 × 10^8^ particles/mL. Zeta potential measurements (–6.0 to –49.0 mV; Fig. 1C) demonstrated surface charge of -18±27.1 mV for carrot EVs, -45.7±3.75 mV for ginger EVs, -30±3.28 mV for garlic EVs, -34±12.4 mV for turmeric EVs indicating stability and indicated minimal aggregation, consistent with the preservation of vesicle integrity [17]. Hence, the obtained edible EVs exhibited intact bilayer structure and optimal purity.

**Figure 1:**
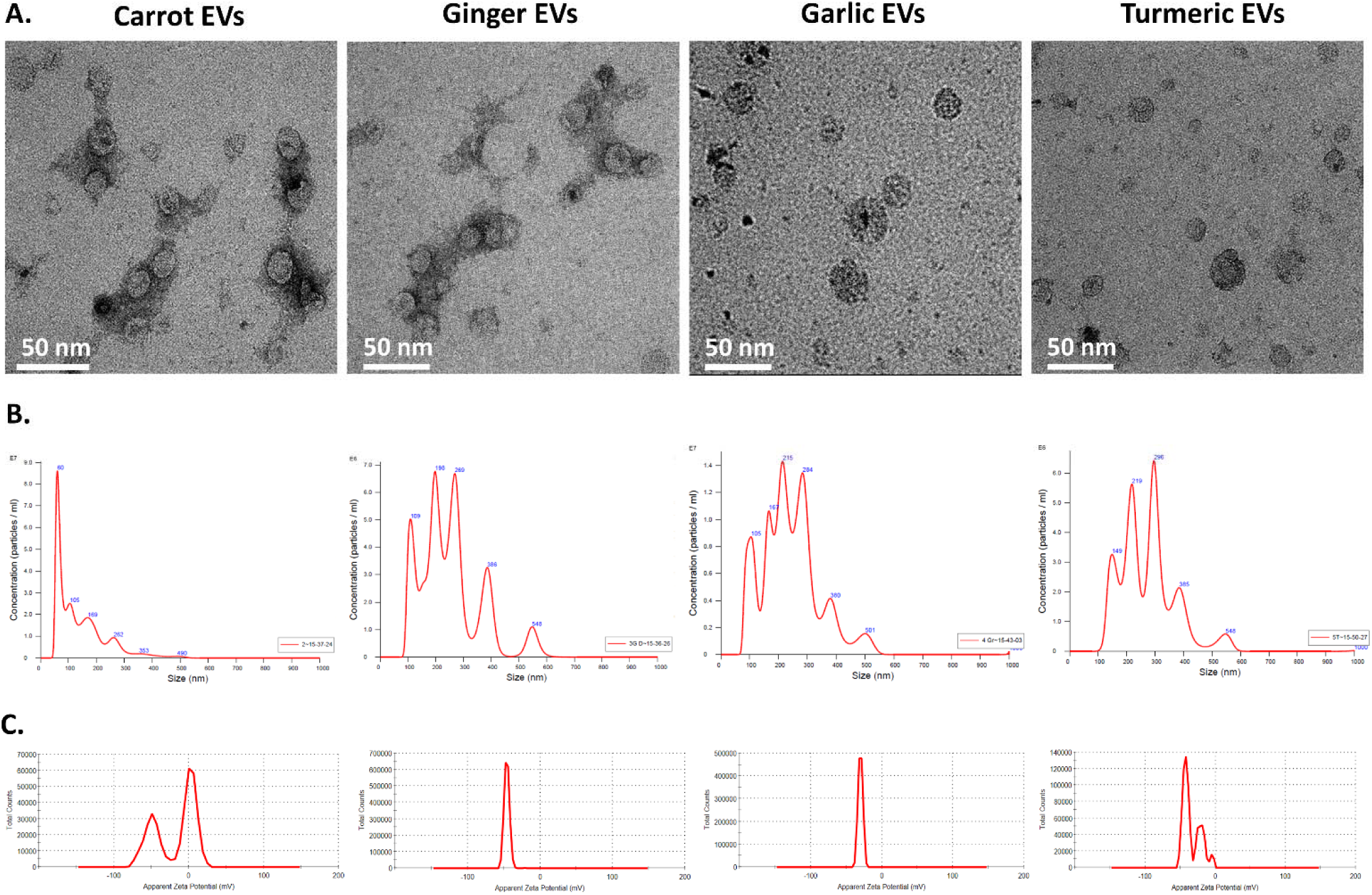
Biological assessment of edible Extracellular Vesicles (EVs). **A.** Morphological identification by Transmission Electron Microscopy (TEM) with scale bar 50nm, **B.** Enumeration of size and concentration of edible EVs using Nanoparticle Tracking Analysis (NTA), **C.** Characterization of edible EVs surface charge by zeta analyzer.

### PDEVs Remain Stable under Gastrointestinal Conditions *In-vitro*

To evaluate the stability of edible EVs under physiologically relevant gastrointestinal conditions, the vesicles were sequentially exposed to simulated SLS followed by ILS, as described in the Methods. NTA analysis demonstrated source-dependent alterations in particle size distribution and concentration following simulated digestion (Fig. 2A). Carrot EVs exhibited a progressive reduction in the predominant mean particle size from 132.3 nm to 74.7 nm after SLS treatment and further to 35.2 nm following ILS digestion, accompanied by an increase in particle concentration from 1.74 × 10^9^ to 5.62 × 10^9^ and 5.54 × 10^9^ particles/mL, respectively. Ginger EVs displayed two predominant particle populations (129.5 and 190.6 nm) following SLS exposure, which converged into a single predominant population at 129.5 nm after ILS treatment, together with a reduction in particle concentration from 3.55 × 10^9^ to 1.14 × 10^9^ and 1.12 × 10^9^ particles/mL, respectively, indicating reduced particle heterogeneity. Garlic EVs exhibited only a marginal reduction in the predominant particle size from 212.5 to 197.9 nm following simulated digestion, with a modest decrease in particle concentration from 2.83 × 10^9^ to 2.41 × 10^9^ particles/mL, suggesting greater physicochemical stability under gastrointestinal conditions. In contrast, turmeric EVs initially exhibited a predominant particle population of 70 nm, which segregated into heterogeneous particle populations of approximately 70, 129, and 189 nm following sequential SLS and ILS digestion, while particle concentration decreased from 2.53 × 10^9^ to 1.55 × 10^9^ particles/mL, indicating preservation of vesicle heterogeneity despite partial particle loss.

**Figure 2:**
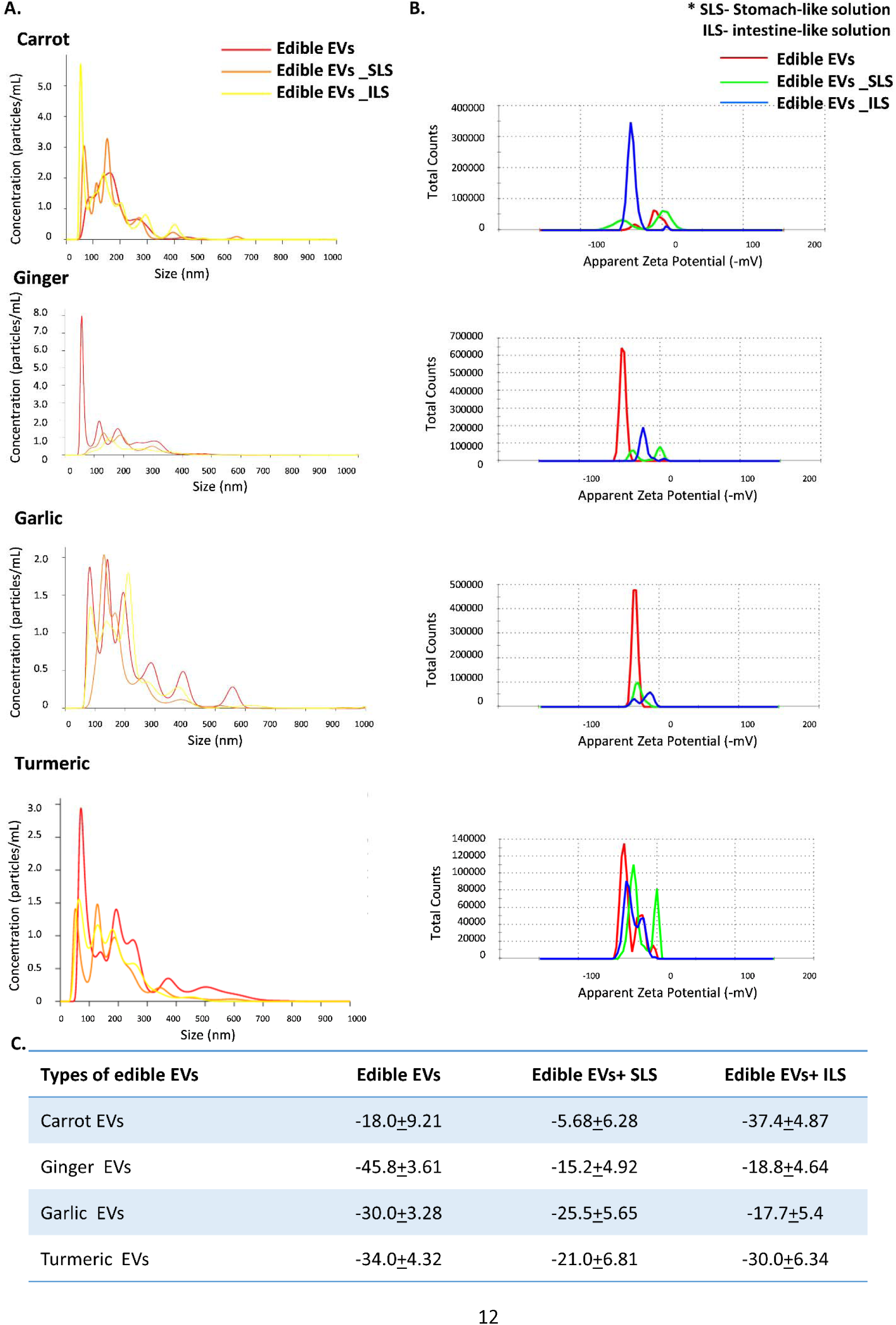
Enumeration and biological stability/viability of edible EVs sourced from carrot, ginger, garlic, and turmeric. **A.** Size distribution and **B.** surface potential of edible EVs, before and after digestion with stomach/intestine-like solutions, **C.** Surface charge of edible EVs before and after digestion with stomach/intestine-like solution. SLS-Stomach-like Solution, ILS-Intestine-like Solution. Values are mean + SD.

Zeta potential analysis (Fig. 2B–C) exhibited source-dependent changes in the surface charge of edible EVs following simulated gastrointestinal digestion. Carrot and turmeric EVs exhibited a similar trend, with an initial reduction in the magnitude of the negative zeta potential following SLS treatment (carrot: −18.0 ± 9.21 to −5.6 ± 6.28 mV; turmeric: −34.0 ± 4.32 to −21.0 ± 6.81 mV), followed by recovery towards their native values after ILS digestion (−37.4 ± 4.87 and −30.0 ± 6.34 mV, respectively). In contrast, ginger and garlic EVs displayed a progressive attenuation of the negative surface charge throughout digestion, with ginger EVs decreasing from −45.8 ± 3.61 to −18.3 ± 4.64 mV and garlic EVs from −30.0 ± 3.28 to −17.7 ± 5.40 mV following sequential SLS and ILS exposure. Nevertheless, all EV preparations retained a net negative surface charge throughout digestion, indicating preserved colloidal stability under simulated gastrointestinal conditions.

### Comprehensive Metabolomic Profiling of PDEVs

A comprehensive metabolomic analysis of all four edible EVs identified 572 annotated compounds, with their sPLSDA distribution illustrated in Fig. 3A. The identified metabolites were distributed across various subcellular locations: 307 in the cytoplasm, 160 in the cell wall, 140 in the membrane, 22 in the ER, 7 in the Golgi apparatus, and 6 in peroxisomes (Fig. 3B). Notably, sphingosine-1-phosphate was enriched in endosomes and glycerophosphocholine was associated with intracellular membranes, suggesting a potential role in endocytic EV biogenesis.

**Figure 3:**
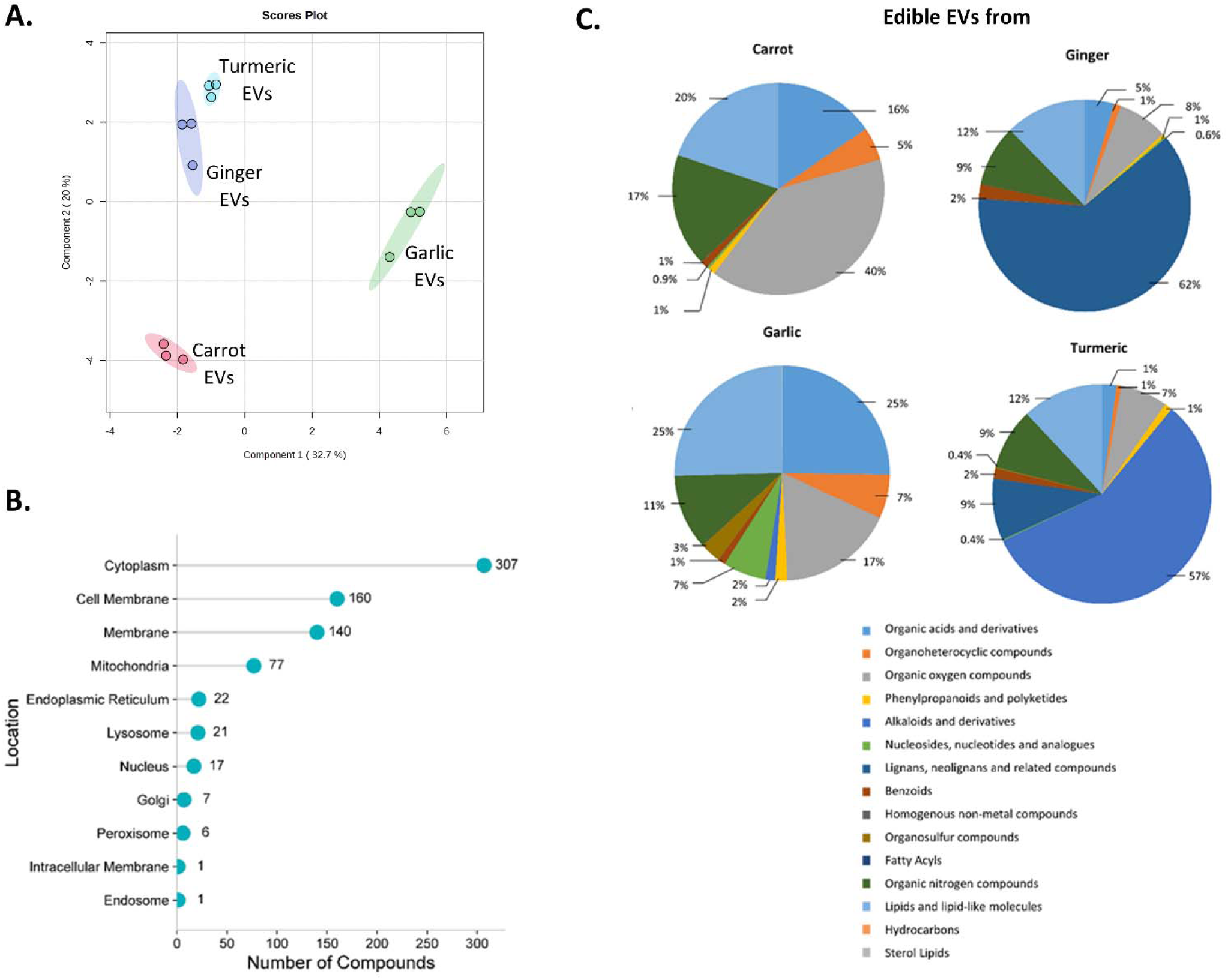
Comprehensive metabolomics of edible EVs. **A.** Sparse Partial Least Squares Discriminant Analysis (sPLSDA) plot depicting metabolome distribution of edible EVs, **B.** Subcellular localization, **C.** Metabolic terrain of edible EVS-associated metabolites.

Compositional analysis further revealed that edible EV-associated metabolites clustered into distinct chemical classes: organic acids and derivatives (OADs), organoheterocyclic compounds (OHC), organic oxygen compounds (OOC), phenylpropanoids and polyketides (PP), alkaloids and derivatives (ADs), nucleosides, nucleotides and analogues (NNA), lignans/neolignans and related compounds (LNC), benzoids, homogenous non-metals (HNM), organosulfur compounds (OSC), fatty acyls, organic nitrogen compounds (ONC), lipids and lipid-like molecules (LLM), hydrocarbons, and sterol lipids (SL). Source-specific distributions were evident. Carrot-derived EVs were enriched in OOC (40%), LLM (20%), ONC (17%), and OAD (16%), with minor contributions from OHC (5%), benzoids (1%), and PP (1%). Ginger EVs were dominated by LNC (62%), with additional contributions from LLM (12%), ONC (9%), OOC (8%), OAD (5%), and minor OHC (1%) and PP (1%). Garlic EVs contained equal proportions of OAD and LLM (25% each), along with OOC (17%), OHC (7%), NNA (7%), OSC (3%), and smaller fractions of PP (2%), AD (2%), and benzoids (1%). Turmeric EVs were primarily composed of AD (57%), with smaller proportions of LLM (12%), LNC and ONC (9% each), OOC (7%), benzoids (2%), and PP (1%), as detailed in Fig. 3C.

### In-Silico Interactome Reveals Source-Specific Association of PDEVs with Distinct Metabolites

Plants are well known to exhibit pharmacological activities attributed to diverse metabolites, including polyphenols, flavonoids, alkaloids, and tannins.[18]. PDEVs are enriched in lipophilic secondary metabolites, highlighting their translational efficacy [19]. Metabolomic analysis identified edible EV-associated metabolites with distinct source-specific abundance patterns across edible extracellular vesicles, as visualized in the heatmap (Fig. 4A). Carrot EVs were found enriched in metabolites such as quercetin, lignin, naringin, pelargonin, xanthotoxin, and quercetin-3-O-glucoside, putatively associated with antioxidant and barrier-protective activities (Suppl.Table S1). Quercetin, lignin and quercetin-3-O-glucoside found strongly interacted with cytochrome P450 enzymes CYP75B1, Glutathione S-transferase F12 GSTF12 (confidence level≥ 0.7; Fig. 4B, Suppl.Table S1), suggesting dietary antioxidant efficacies. Ginger EVs are enriched with xanthine, silymarin, (-)-epicatechin, sinapyl alcohol, and [6]-gingerol (Suppl. Fig. S2B). [6]-Gingerol and silymarin show strong interactions with tumor protein (TP53), cyclin D1 (CCND1), matrix metalloproteinases (MMP9), while linking to catalase (CAT) and CYP3A4 (Fig. 4.C, Suppl. Table S2). Xanthine interacted with Purine Nucleoside Phosphorylase (PNP), Guanine Deaminase (GDA), and Hypoxanthine-Guanine Phosphoribosyltransferase (HPRT1).

**Figure 4.**
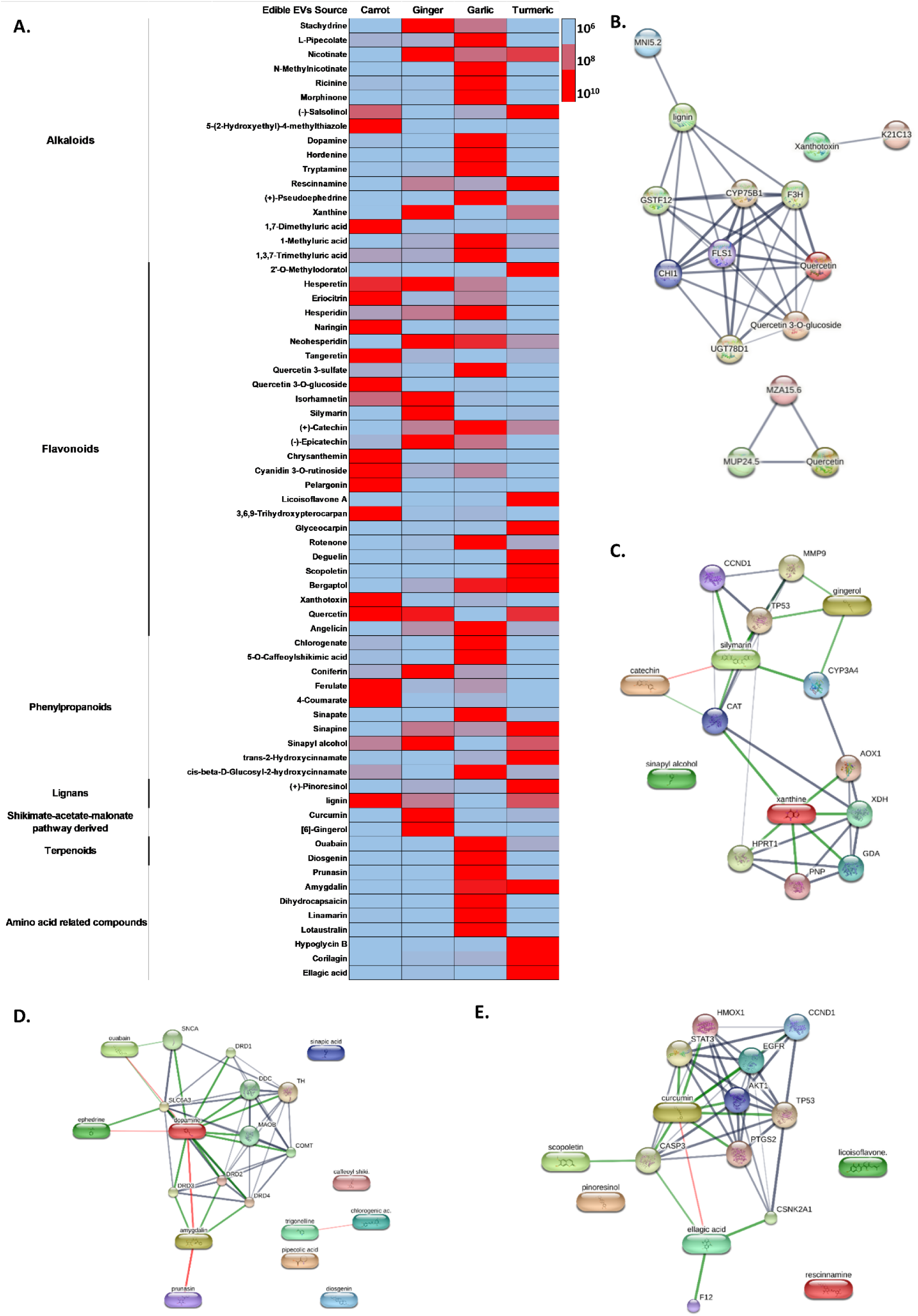
Metabolites in edible Extracellular Vesicles (EVs) and their predicted interactomes. **A.** Distribution of key phytocompounds identified in edible EVs, predicted interactome networks of phytocompounds from edible EVs derived from **B.** carrot, **C.** ginger, **D.** garlic, **E.** turmeric, highlighting source-specific molecular targets and potential therapeutic pathways. Line thickness indicates the strength of data support. Red line indicates the presence of fusion evidence. Green line indicates neighborhood evidence. Gray line indicates coexpression evidence.

Garlic EVs contained bioactives such as L-pipecolate, N-methylnicotinate, (+)-pseudoephedrine, chlorogenate, 5-O-caffeoylshikimic acid, sinapate, diosgenin, dopamine, ouabain, prunasin, and amygdalin (Suppl. Fig. S2.C). Dopamine, ouabain, and amygdalin were found associated with monoamine oxidase (MAO-B), dopamine receptors (DRD1-4) and transporter (SLC6A3), tyrosine hydroxylase (TH), and amyloid precursor protein synuclein (SNCA) in Fig. 4D and Suppl.Table S3. Diosgenin, enriched in garlic EVs, is known for its anti-cholesterolemic effects [20], and chlorogenate exhibited antisteatotic activity [21]. Turmeric EVs carry rescinnamine, scopoletin, (+)-pinoresinol, licoisoflavone A, ellagic acid, and curcumin (Suppl. Fig. S2D, Table S4). Curcumin, the principal bioactive (Fig. 4E), showed strong interactions with epidermal growth factor receptor (EGFR), RAC-alpha serine/threonine-protein kinase (AKT1), signal transducer and activator of transcription 3 (STAT3), prostaglandin-endoperoxide synthase 2 (PTGS2), TP53, and CASP3. Additionally, ellagic acid and scopoletin demonstrated strong associations with CASP3.

### Edible Extracellular Vesicles Restore Gut Epithelia *In Vitro*

Annotated interactome analysis suggested that edible EV-associated metabolites participate in the regulation of antioxidant responses, cell proliferation, cell cycle progression, apoptosis, and related signaling pathways. Given their intended oral administration, the biological functionality of edible EVs was evaluated using HCT-116 cells as an *in vitro* intestinal epithelial model (schematic in Fig. 5A). Cellular stress was induced using 20 mM NH₄Cl, followed by treatment with edible EVs isolated from the five plant sources. Subsequently, the mRNA expression of genes associated with epithelial barrier integrity, inflammation, antioxidant defense, cell proliferation, cell cycle regulation, and apoptosis was evaluated.

**Figure 5:**
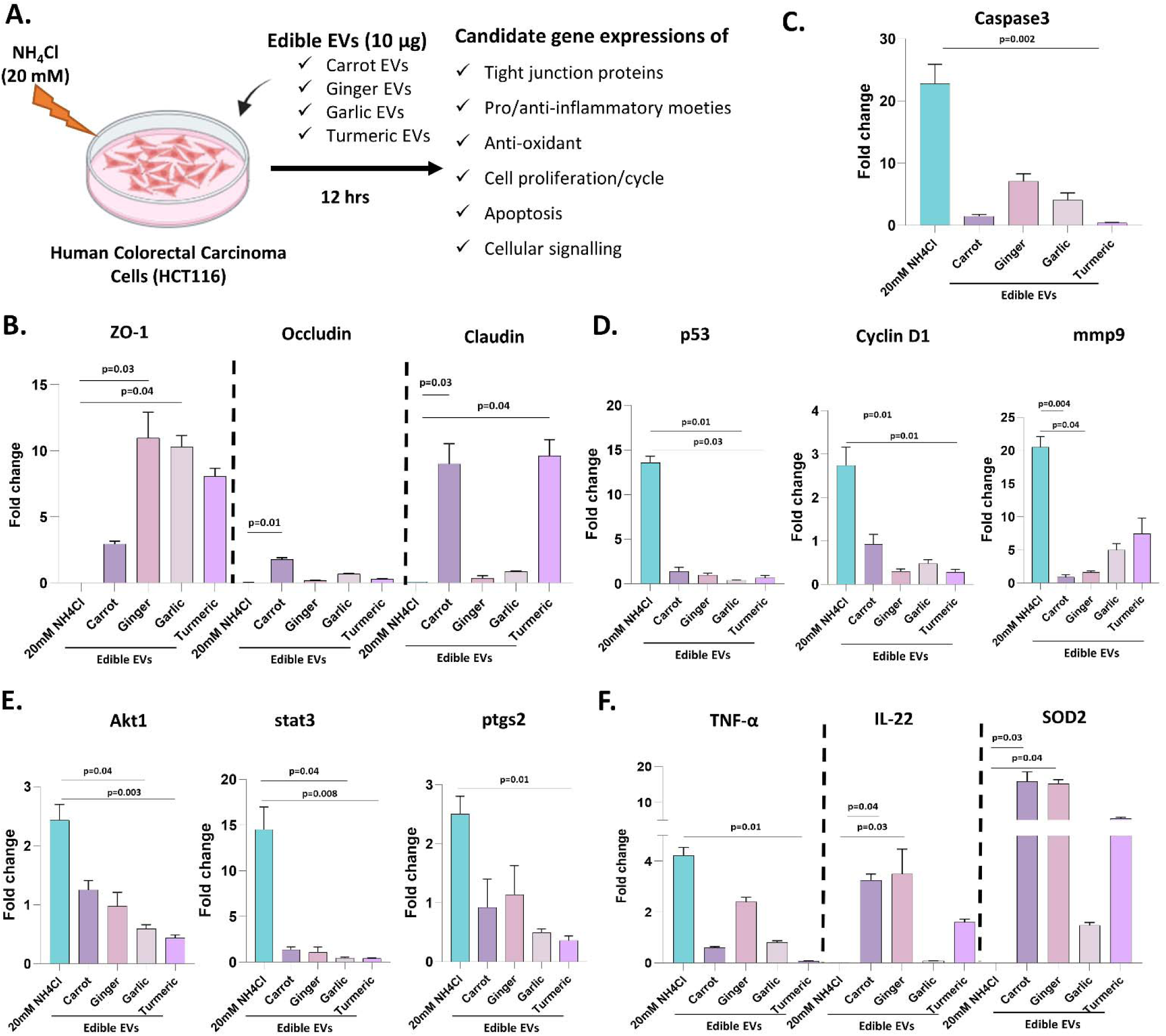
*In vitro* Pharmacological assessment of edible EVs. **A.** Illustration of the *In Vitro* Co-Culture System Simulating Enterocyte Stress and edible EVs treatment. Gene expressions of **B.** Junctional communication proteins and **C.** apoptosis, **D.** Cell proliferation/cycle, **E.** Cellular signalling, **F.** pro-inflammatory, anti-inflammatory and anti-oxidative genes post Edible EVs treatment.

Edible EVs from different sources modulated the expression of tight junction-associated genes in NH₄Cl-stimulated HCT-116 cells (Fig. 5B). Restoration of ZO-1 expression was observed following treatment with carrot (>2.5-fold), ginger (>10-fold), garlic (>8-fold), and turmeric (>8-fold) EVs, with ginger and garlic EVs exhibiting the most pronounced effects (p < 0.05). In contrast, carrot EVs specifically enhanced the expression of occludin (>2-fold, p < 0.05) and claudin (>8-fold, p < 0.05), whereas turmeric EVs also significantly upregulated claudin expression compared with the NH₄Cl-treated group (>9-fold, p < 0.05).

Turmeric EVs significantly reduced caspase 3 (>20-fold, p = 0.002; Fig. 5C). Garlic and turmeric EVs significantly downregulated p53 (>12-fold, p< 0.05). Ginger and turmeric EVs downregulated cyclin D1 (>2.5-fold, p< 0.05, fig. 5D). Carrot and ginger EVs downregulated mmp9 (>20-fold, p< 0.05). Garlic and turmeric EVs significantly downregulated akt1 (>2.5-fold, p < 0.05) and stat3 (>15-fold, p < 0.05; Fig. 5E). Whereas, turmeric EVs specifically reduced the expressions of ptgs2 (>2-fold, p < 0.05; Fig. 5E). Consistently, edible EVs treatment markedly downregulated TNF-α, particularly with turmeric EVs (>4-fold; p = 0.01, Fig. 5F) and upregulated IL-22 with carrot EVs (>3.5-fold; p = 0.002); ginger EVs (>1.5-fold; p = 0.01) compared to NH₄Cl stimulation. Fig. 5C shows enhanced expression of SOD2 with carrot (p = 0.03), and ginger (p = 0.04) demonstrating restoration of epithelial integrity with anti-inflammatory and antioxidant efficacy.

### Edible Extracellular Vesicles Exert Hepatoprotective Effect *in vitro*

Although in silico analyses predicted the hepatoprotective potential of edible EV-associated metabolites, their ability to reach and interact with hepatic cells remains to be experimentally validated. Therefore, the hepatic uptake of edible EVs was subsequently investigated. Flow cytometry analysis (Fig. 6A) revealed that carrot and turmeric EVs exhibited the highest uptake by HepG2 cells compared to ginger and garlic EVs. Z-stack confocal imaging (Fig. 6B) further confirmed the internalization of PKH26-labeled carrot EVs (red), which were visualized at a depth of 3 µm within IHH cells but not at the surface. PKH26-labeled carrot EVs accumulated at the hepatocyte membrane during the process of internalization. Live-cell confocal microscopy captured dynamic uptake of PKH26-labeled carrot EVs in IHH cells, with images recorded every 5 seconds over a 20-minute period (Suppl. Fig. S3A). Quantitative analysis showed maximal internalization within 15 min of co-culture, with fluorescence intensity measurements of labeled edible EVs depicted in Suppl. Fig. S3B.

**Figure 6:**
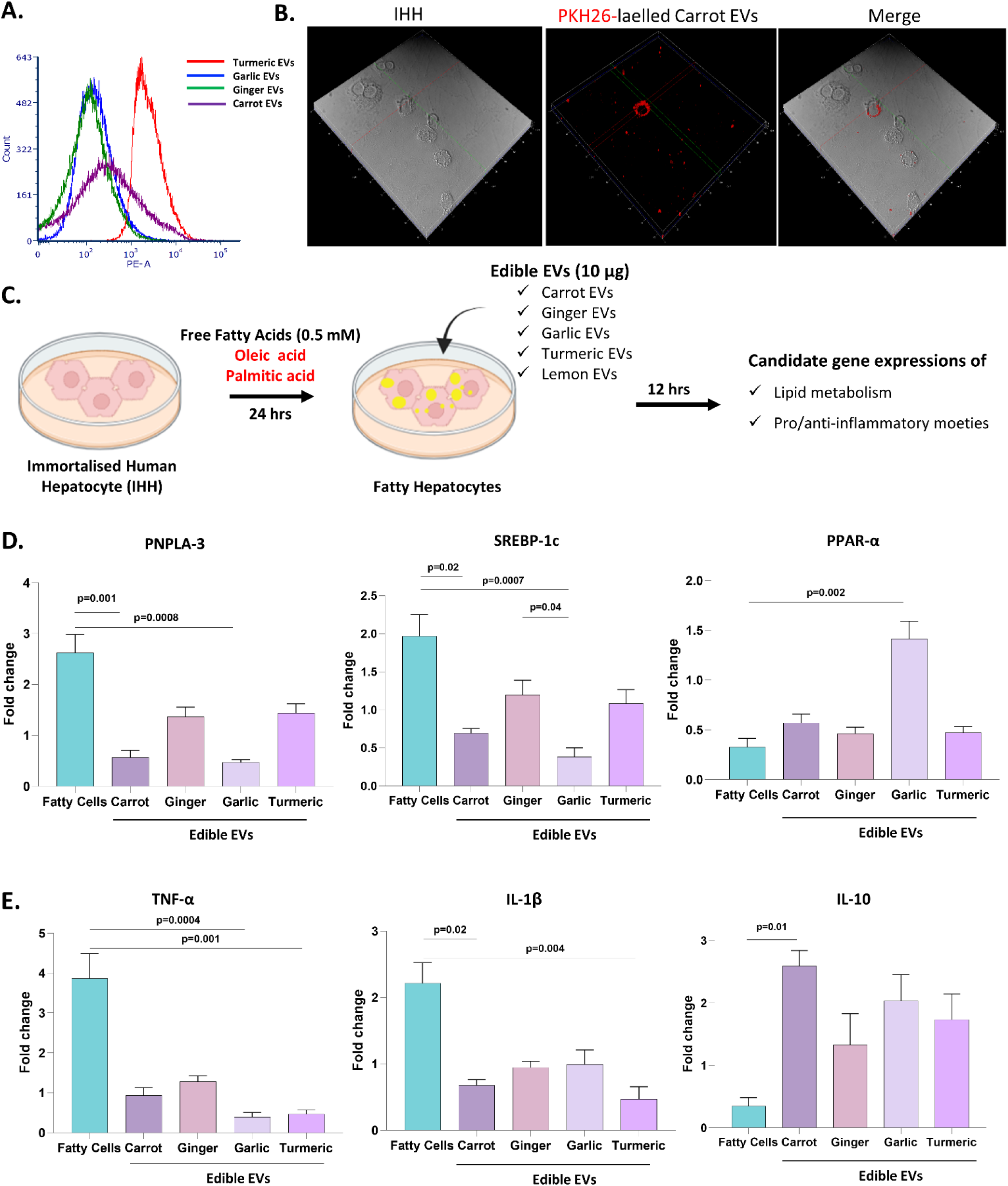
Hepatic uptake of PKH26-labelled edible EVs *in vitro*, Relative internalization of PKH26-labelled edible EVs **A.** by HepG2 cells analysed using Flow cytometry, **B.** Z-stack confocal microscopy analysis of PKH26-labelled carrot EVs by primary hepatocyte IHH, **C.** Illustration of the *in vitro* fatty cell culture model, Simulation with 0.5 mM free fatty acids (oleic and palmitic acids) to immortalised human hepatocytes (IHH) and edible EVs treatment. Expressions of genes associated to **D.** Lipid metabolism and **E.** pro-inflammation, and anti-inflammation post-edible EVs treatment. Statistical test used Kruskal-Wallis, p-values compared to NH₄Cl stimulation and fatty IHH cells developed with free fatty acid stimulation respectively.

To further evaluate the hepatic functionality of edible EVs, their effects were assessed in an *in vitro* steatosis model. Steatosis was induced in IHH using 0.5 mM free fatty acids (FFA; oleic acid:palmitic acid) (Fig. 6C), resulting in significant alteration of genes associated with lipid metabolism and inflammation. Treatment with edible EVs restored the expression of these genes. Specifically, carrot- and garlic EVs significantly downregulated the expression of PNPLA3 (>2.5-fold, p < 0.001; Fig. 6D) and SREBP-1c (>1.5-fold, p < 0.05). In addition, garlic EVs significantly upregulated PPAR-α expression (>1.5-fold, p = 0.002), indicating improved regulation of hepatic lipid metabolism. Notably, garlic EVs produced a significantly greater reduction in SREBP-1c expression than ginger EVs (p = 0.04). Edible EVs differentially modulated the inflammatory response in NH₄Cl-stimulated hepatocytes by suppressing pro-inflammatory cytokines while promoting anti-inflammatory signaling (Fig. 6E). Garlic- and turmeric-derived EVs significantly downregulated TNF-α expression by >4.5-fold (p < 0.001), whereas carrot- and turmeric-derived EVs markedly reduced IL-1β expression (>1.5-fold, p < 0.05). In contrast, carrot EVs significantly enhanced the expression of the anti-inflammatory cytokine IL-10 (>2.5-fold, p < 0.05), indicating their immunomodulatory potential.

## Discussion

The present study demonstrates that edible plant-derived extracellular vesicles (PDEVs) isolated from four phytochemically distinct food sources possess unique metabolite signatures that are associated with distinct functional properties. By integrating physicochemical characterization, simulated gastrointestinal digestion, untargeted metabolomics and functional validation, this study provides one of the first comparative evaluations that evaluated metabolites, linking food-derived secondary metabolites packaged within naturally occurring nanovesicles to source-specific biological activities. These findings extend current knowledge by suggesting that edible PDEVs may represent naturally occurring food nanostructures that preserve and deliver phytochemicals while contributing to the functional properties of edible plants.

The persistence of PDEVs following simulated gastrointestinal digestion represents an important finding from a food science perspective. The nutritional efficacy of many plant-derived secondary metabolites is often constrained by poor gastrointestinal stability, extensive metabolism, and limited bioaccessibility after oral consumption. Although digestion induced source-dependent changes in particle size distribution and surface charge, vesicular integrity was largely retained, suggesting that the lipid bilayer undergoes physicochemical remodeling rather than complete disruption under gastrointestinal conditions. Such membrane protection may enhance the stability of metabolites and facilitate their interaction with intestinal epithelial cells, thereby improving their functional availability. Similar observations have been reported for ginger-derived extracellular vesicles, which remain stable during gastric digestion and retain biological activity following oral administration, supporting the concept that edible plant vesicles naturally protect bioactive constituents during gastrointestinal transit [6, 8]

A major outcome of this study is the demonstration that the metabolite composition of PDEVs is highly dependent on the botanical source. Untargeted metabolomics representing diverse classes of food bioactives, including flavonoids, phenylpropanoids, terpenoids, alkaloids, sterol lipids, and fatty acyls. Metabolomic profiling of edible EVs showed a diverse repertoire of organic acids, phenylpropanoids, polyketides, alkaloids, fatty acyls, sterol lipids, and other bioactive classes. Whereas, subcellular annotation identified metabolites associated with the plasma membrane, endoplasmic reticulum, Golgi apparatus, endosomes, intracellular membranes, and peroxisomes, providing complementary evidence consistent with an endomembrane-derived vesicular origin. Nevertheless, future studies incorporating orthogonal EV purification strategies, as recommended by the MISEV guidelines [14], will further strengthen the vesicle-specific interpretation of the observed biological and metabolomic findings.

*In silico* analyses further connected these metabolites to key signaling pathways, providing mechanistic insights into their therapeutic actions. Carrot EVs, enriched with quercetin, lignin, and quercetin-3-O-glucoside, exhibited strong interactions with CYP protein and glutathion transferase indicative of pronounced antioxidative potential. Furthermore, carrot EVs suppressed inflammation while upregulating IL-22, IL-10 and SOD2, reinforcing their barrier-protective, anti-inflammatory and antioxidant functions in alignment with previous findings [11]. Similarly, ginger EVs were enriched with 6-gingerol and silymarin, which were associated with key regulators of cell-cycle progression. Consistent with these interactions, ginger EVs downregulated Cyclin D1 and MMP9 expression in NH₄Cl-stressed HCT-116 cells, suggesting suppression of ammonia-induced proliferative and invasive signaling. Xanthine enrichment further indicated a potential role in nucleotide metabolism through its association with HPRT1, PNP, GDA, and XDH. Moreover, the upregulation of IL-22 highlights the anti-inflammatory potential of ginger EVs, consistent with the protective effects reported by Teng et al. in experimental colitis [6].

Garlic EVs were enriched in diosgenin and chlorogenate, phytometabolites previously reported to possess anti-cholesterolemic [20] and anti-steatotic [21] activities. Consistent with these reports, garlic EVs modulated lipid metabolism in fatty acid-induced hepatocytes by suppressing PNPLA-3 and SREBP-1c expression while enhancing PPAR-α-associated fatty acid oxidation. These findings complement Liu *et al.*, who identified miR-369e in garlic EVs as a key regulator of lipid metabolism via macrophage–hepatocyte crosstalk in fatty acid-fed mice [10]. Garlic EVs also reduced TNF-α in hepatocytes, and in colon epithelial cells, while enhancing ZO-1 expression, supporting the anti-inflammatory effects reported by Zhao *et al.* [22]. Moreover, dopamine and amygdalin suggest potential modulation of dopamine homeostasis through TH, MAO-B, DRD1–4, SNCA, and SLC6A3, though molecular validation is required in future studies.

Turmeric EVs, enriched with curcumin and scopoletin, mitigated stress-induced colon epithelial damage by modulating the EGFR–AKT1/STAT3–PTGS2 axis and restoring the survival–apoptosis balance. They suppressed p53, TNF-α, and PTGS2, reduced IL-1β, and enhanced IL-10 and lipid homeostasis in hepatocytes, demonstrating potent anti-inflammatory and hepatoprotective activity. These effects align with previous LPS-induced and DSS-induced colitis mice models [13, 23], suggesting these molecular pathways underlie the anti-inflammatory mediated mechanism of turmeric EVs. Rapid hepatic uptake of PKH26-labeled carrot, turmeric, and garlic EVs within 15 minutes reinforced their translational relevance, consistent with previous observations [24].

In conclusion, edible PDEVs retain source-specific associated metabolites with distinct functional properties and remain stable under simulated gastrointestinal conditions. These naturally occurring food nanocarriers represent promising functional food ingredients that may enhance the bio-accessibility and biological efficacy of dietary phytochemicals for promoting gut and liver health.

## Acknowledgements

We extend our gratitude to Dr B.S Tomar, Indian Agricultural Research Institute (IARI) and the National Horticultural Research and Development Foundation (NHRDF), New Delhi, and Kasam, Odisha for generously providing the edible plant species used in this study. We acknowledge ICMR (Indian Council of Medical Research) New Delhi for funding the fellowship to Ms. P. Debishree Subudhi.

## Data Availability Statement

Raw data of metabolomics have been provided as supplementary and other Data will be made available on request.

## Conflict of Interest Statement

All authors have declared no conflict of interest.

## Financial Support

This work was supported by SERB Power Grant no. SPG/2021/002862

## Declaration of Generative AI

During the writing of the manuscript, authors used ChatGPT (OpenAI) in order to improve the language. After using this tool, the authors reviewed and edited the content as needed and take full responsibility for the content of the publication.

## List of Abbreviations

ACN: Acetonitrile
ADs: Alkaloids and derivatives
AKT1: Alpha serine/threonine-protein kinase 1
CAT: Catalase
CCND1: Cyclin D1
CYP: Cytochrome P450 enzymes
DRD1-4: Dopamine receptors 1-4
DUC: Differential ultracentrifugation
Edible EVs: Edible extracellular vesicles
EGFR: Epidermal growth factor receptor
ER: Endoplasmic reticulum
GDA: Guanine deaminase
GUC: Sucrose gradient ultracentrifugation
HCAR1-3: Hydroxycarboxylic acid receptors 1-3
HNM: Homogenous non-metals
HPRT1: Hypoxanthine-guanine phosphoribosyltransferase 1
IHH: Immortalized human hepatocyte
IL: Interleukin
ILS: Intestine-like solution
LC-MS: Liquid chromatography–mass spectrometry
LLM: Lipids and lipid-like molecules
LNC: Lignans, neolignans, and related compounds
MAO-B: Monoamine oxidase B
MMP9: Matrix metalloproteinase 9
NAPRT1: Nicotinate phosphoribosyltransferase 1
NH₄Cl: Ammonium chloride
NNA: Nucleosides, nucleotides and analogues
NR1I2: Nuclear receptor subfamily 1 group I member 2
NTA: Nanoparticle tracking assay
OADs: Organic acids and derivatives
OHC: Organoheterocyclic compounds
ONC: Organic nitrogen compounds
OOC: Organic oxygen compounds
OSC: Organosulfur compounds
PEVs: Plant-derived extracellular vesicles
PLSDA: Partial least squares discriminant analysis
PNP: Purine nucleoside phosphorylase
PNPLA3: Patatin-like phospholipase domain-containing protein 3
PP: Phenylpropanoids and polyketides
PPAR-α: Peroxisome proliferator-activated receptor alpha
PTGS2: Prostaglandin-endoperoxide synthase 2
qRT-PCR: Quantitative real-time polymerase chain reaction
SL: Sterol lipids
SLS: Stomach-like solution
SNCA: Synuclein alpha (amyloid precursor)
SOD: Superoxide dismutase
SREBP-1c: Sterol regulatory element-binding protein 1c
STAT3: Signal transducer and activator of transcription 3
TUB: Tubulins
TNF-α: Tumor necrosis factor alpha
TOP: Topoisomerase
TP53: Tumor protein p53
ZO-1: Zonula occludens-1

## Notes

### Competing Interest Statement

The authors have declared no competing interest.

